# The Evolutionary Moulding in plant-microbial symbiosis: matching population diversity of rhizobial *nod*A and legume *NFR5* genes

**DOI:** 10.1101/285882

**Authors:** Anna A. Igolkina, Georgii A. Bazykin, Elena P. Chizhevskaya, Nikolai A. Provorov, Evgeny E. Andronov

## Abstract

We propose the Evolutionary Moulding hypothesis that population diversities of partners in nitrogen-fixing rhizobium-legume symbiosis are matched, and tested it in nucleotide polymorphism of symbiotic genes encoding two components of the plant-bacteria signalling system. The first component is the rhizobial *nod*A acyltransferase involved in the fatty acid tail decoration of Nod factor (rhizobia signalling molecule). The second component is the plant *NFR5* receptor, putatively required for Nod-factor binding.

We collected three wild growing legume species together with soil samples adjacent to the roots (soil pool) from one large 25-year fallow: *Vicia sativa, Lathyrus pratensis* and *Trifolium hybridum* nodulated by one of the two *Rhizobium leguminosarum* biovars (*viciae* and *trifolii*). For each plant species we prepared three pools for DNA extraction: the plant pool (30 plant indiv.), the nodule pool (90 nodules) and the soil pool (30 samples). *NFR5* gene libraries from the plant pool and *nod*A gene libraries from nodule and soil pools were sequenced by Sanger technology and High-throughput pyrosequencing, respectively. Analysis of the data demonstrated concordance in population diversities of one symbiotic partner (rhizobia) the second partner (legume host), in line with the Evolutionary Moulding hypothesis. This effect was evinced by the following observations for each plant species: (1) significantly increased diversity in the nodule *nod*A popset (set of gene sequences derived from the nodule population) compared to the soil popset; (2) a monotonic relationship between the diversity in the plant *NFR5* gene popset and the nodule rhizobial *nod*A gene popset; and (3) higher topological similarity of the *NFR5* gene tree with the *nod*A gene tree of the nodule popset, than with the *nod*A gene tree of the soil popset. Both nonsynonymous diversity and Tajima’s D were increased in the nodule popsets compared to the soil popsets, consistent with relaxation of negative selection and/or admixture of balancing selection underlying the Evolutionary Moulding effect. We propose that the observed genetic concordance arises from the selection of particular characteristics of the nodule *nod*A genes by the host plant.

## Introduction

One of the earliest studied types of symbiosis, the host–parasite interaction, was described by Flor’s Gene-for-Gene concept (Flor 1971; Flor 1942) and, in fact, the first mathematical model of coevolution was explicitly based on the assumption of a Gene-for-Gene (GFG) interaction (Thompson and Burdin 1992; Mode 1958). Further analyses of host-parasite interactions revealed alternative concepts, namely matching-allele(MA) (Frank 1994), inverse-matching-allele (IMA) (Otto and Michalakis 1998) and inverse-gene-for-gene (IGFG) (Fenton, Antonovics, and Brockhurst 2009), which together with the GFG represent the opposite end of the same continuum of host–parasite specificity (Agrawal and Lively 2002). These theoretical concepts, developed for antagonistic systems, found their reflection in mutualistic symbiotic systems (Lewis-Henderson and Djordjevic 1991; Sadowsky et al. 1991; Cregan, Sadowsky, and Keyser 1991; Parker 1999; Sachs, Essenberg, and Turcotte 2011). Later, various mathematical and theoretical models were developed to describe different types of symbiotic interactions (Yoder 2016; Kwiatkowski, Engelstadter, and Vorburger 2012). One aspect of symbiotic interactions that previously had much attention in models is the difference in evolutionary rates between interacting species. For example, the so-called Red Queen model (Van Valen 1973; Paterson et al. 2010; Pal et al. 2007) suggests the metaphor of an evolutionary arms race, whereby a more rapidly evolving species in a host–parasite interaction has an adaptive advantage over a more slowly evolving species. For mutualistic symbiosis, the alternative Red King model proposes that the slower-evolving species are favoured (Bergstrom and Lachmann 2003). However, the recent study showed that the preference of the particular model is not uniquely determined by the type of symbiosis (Rubin and Moreau 2016). Specifically, the Red Queen model initially introduced for antagonism was detected in the ant-plant mutualism. Here, we suggested another aspect of symbiotic interactions that can potentially provide the basis for models describing both mutualism and antagonism.

Our hypothesis, which we refer to as Evolutionary Moulding, is that the population diversity levels and structures are matched between symbionts. We proposed that this hypothesis can be a common aspect of symbiotic systems and tested it in the nitrogen-fixing rhizobium-legume symbiosis. Previous analysis of the symbiosis between *Rhizobium galegae* and its host plant *Galega* indicated correspondence of population diversity levels between microsymbionts and the host *Galega* species (Andronov et al. 2003; Österman et al. 2011). In particular, a more genetically diverse *Galega orientalis* population harbours a more diverse root-nodule rhizobial population, while its less diverse sympatric counterpart *Galega officinalis* forms symbiosis with a less diverse rhizobial population. This observation is related to the well-studied phenomenon of shaping the genetic structure of the rhizobial population through the selection of specific rhizobial genotypes by the host plant (Paffetti et al. 1998; Laguerre et al. 2003; Heath and Tiffin 2007; Depret and Laguerre 2008). The reverse effect, i.e. plant evolution driven by rhizobia, has also been discussed (Martínez-Romero 2009), and is consistent with the recently detected evidence for balancing selection on the symbiotic legume genes in a large legume genome dataset (Yoder 2016). Therefore, we expected that the interplay of symbiotic populations leads to concordance between the diversity levels in their symbiotic genes.

In our study, we focused on two genes encoding the essential components of rhizobium-legume signalling system which are associated with each other through a lipochito-oligosaccharide called Nod factor (Figure 1). The first component is the rhizobial *nod*A gene which encodes an acyltransferase enzyme essential in Nod factor (NF) biosynthesis, specifically in the attachment of the long-chain fatty acid tail to the oligosaccharide backbone (Dénarié, Debellé, and Promé 1996; Esseling and Emons 2004; Oldroyd 2013). The second component is one of the plant symbiotic receptor genes, *NFR5*, which is a homologue of *LjNFR5, MtNFP, PsSym10* genes. Its product recognises NFs (signalling molecules) by three extracellular LysM domains and triggers the formation of root nodule primordia giving the green light to the process of bacterial infection (Oldroyd 2013). The NFs are major determinants of host specificity: rhizobia produce NFs with different structures and host plants percept only those Nod factors that have a certain composition (Mergaert and Montagu 1997). The variation of NFs structure is observed not only between rhizobia species but also at the intra-species level (Spaink 2000), one rhizobia species produces a mixture of NFs that vary in the fatty tail modifications. As proposed, the *nod*A product can vary in its fatty acid specificity, thus contributing to the bacterial host range (Moulin, Béna, and Stępkowski 2004; Dénarié, Debellé, and Promé 1996; Ritsema et al. 1996; Roche et al. 1996). It is logical to assume that the *nod*A gene diversity in a rhizobial population can reflect the structural variation of NFs produces by this population. Indeed, it has been shown that minor differences in the structure of fatty acids tail can affect intra-species host specificity (Li et al. 2011). On the host-plant side, NFs are recognized by high-affinity legume receptors (Broghammer et al. 2012; Moulin, Béna, and Stępkowski 2004). Studies on the model legumes revealed *NFR5* as one of the major receptors to percept NFs (Radutoiu et al. 2007). Mutant analysis showed that single amino acid differences in one domain of the *NFR5* receptor change recognition of Nod factor variants (Broghammer et al. 2012; Radutoiu et al. 2007). Such mediation of NFs between rhizobial *nod*A and legume *NFR5* genes make them good candidates for Evolutionary Moulding.

**Figure 1.**
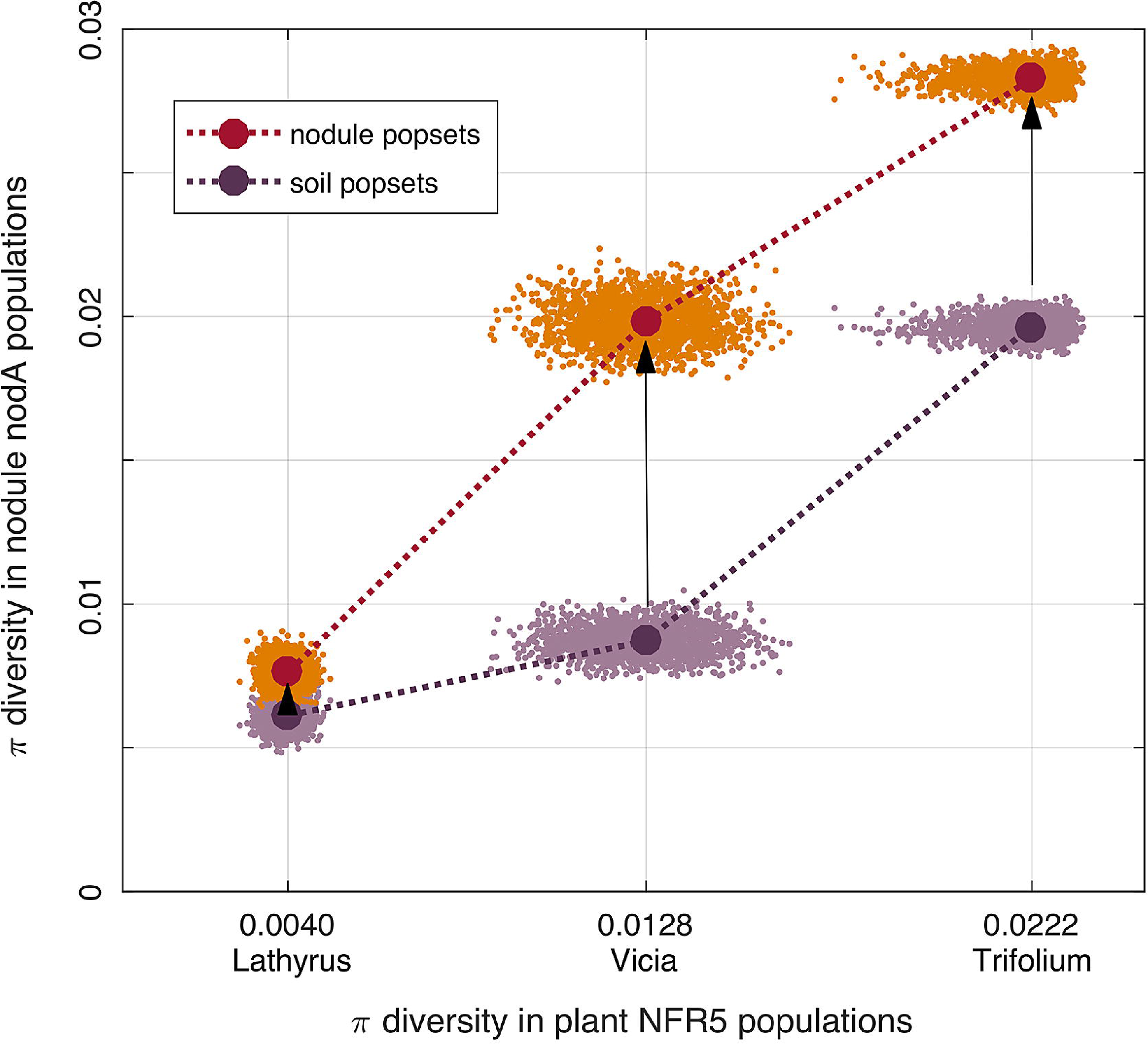
A part of the signal transduction system that governs the rhizobium-legume symbiosis. The rhizobial *nod*A gene encodes the acyltransferase that participates in the attachment of the hydrophobic long-chain fatty acid tail to the Nod factor core. Plant *NFR5* gene encodes the symbiotic receptor recognising the rhizobial Nod factor followed by the symbiosis formation.

Moreover, it has been shown that the topology of the *nod*A gene tree follows the corresponding host plant tree more strictly than the 16S rRNA-based rhizobial phylogeny (Dobert, Breil, and Triplett 1994; Suominen et al. 2016). Thus, we proposed that the Evolutionary Moulding can be observed by comparing the genetic diversity levels of the symbiotic genes of both partners.

We investigated the *nod*A-*NFR5* symbiotic systems of three wild growing legume species (*Vicia sativa, Lathyrus pratensis* and *Trifolium hybridum*) and their rhizobial microsymbionts. Sampling of the experimental material was performed uniformly on the one large natural fallow (more than 25 years) field in order to avoid the influence of ecological factors. *Vicia* and *Lathyrus* species represent the same cross-inoculation group nodulated with *Rhizobium leguminosarum* bv. *viciae* strains, while *Trifolium* belongs to a separate group nodulated with *R.leguminosarum* bv. *trifolii*. One of the important traits of the rhizobium-legume symbiosis is the annual cycle of rhizobia, consisting of nodule formation with consequent amplification of rhizobia inside of the nodule, followed by a release of the nodule rhizobia back into the soil after nodule degradation probably leading to an increase of the frequency of this rhizobial genotype in the soil. Therefore, we analysed both soil and nodule populations of rhizobia, which affect each other.

Testing the Evolutionary Moulding hypothesis required comparison of structural (topological) characteristics of plant and rhizobia populations. Traditionally, the topological similarity between two populations is estimated as the congruence of two respective labelled trees (Leigh et al. 2011). Here, we proposed a novel method to compare topologies of two gene trees with unlabelled leaves. The method was based on the gCEED approach (Choi and Gomez 2009) that translates each population to the Gaussian mixture model in a K-dimensional space. This method can be classified as a kind of beta-diversity metrics, which, by analogy with taxonomic (Jost, Chao, and Chazdon 2011) and phylogenetic (e.g. UniFrac (Lozupone et al. 2011)), could be denoted as “topological beta-diversity”. We applied it to show that the tree structures are concordant between the two symbiont species.

The main aim of this study was to estimate and to compare the population diversity of the symbiotic genes between the soil and nodule rhizobial populations and to link the observed differences to those between the symbiotic determinants of the host plants. Using common diversity statistics and a novel approach for comparing the structures of partner’s populations from pooled sequencing data we confirmed the Evolutionary Moulding hypothesis.

## Materials and Methods

### Sampling

Three wild growing legume species (30 samples per species) together with rhizosphere soil-the common vetch *Vicia sativa*, the meadow vetchling *Lathyrus pratensis* and the alsike clover *Trifolium hybridum* – were uniformly collected from the large natural fallow (more than 25 years) field near the town Vyritsa (Gatchinskii region of Leningradskaya oblast, Russia, 59°24′7.74′′ N; 30°15′28.74′′ E). All sampled plants had formed nitrogen-fixing symbiotic root nodules which were selected and thoroughly washed. Soil samples were collected from the close proximity to the plant roots (1-5 cm). Here and below we named these samples as “soil samples”. Three nodules from each plant sample were picked from the main or the closest to the main lateral roots. For each legume species we prepared three pools for DNA extraction: the plant pool (30 leaf pieces, 0.1g each), the nodule pool (90 nodules, 3 nodules per plant individual) and the soil pool (30 soil samples, 0.2g each).

### *nod*A amplification and sequencing

DNA was isolated from soil and nodule pools by bead beating homogenization (Precellys 24) and purification (PowerSoil DNA Isolation Kit, MoBio, United States). Two pairs of nested degenerate oligonucleotide primers were designed for *nod*A gene of *R.leguminosarum* bv. *viciae* and bv. *trifolii*. The first round of nested PCR with external primers – forward (5’-DGGHYTGTAYGGAGTGC-3’) and reverse (5’-AGYTCSSACCCRTTT −3’) – produces a 324bp amplicon product; the second round with inner primers – forward (5’-YTDGGMATCGCHCACT-3’) and reverse (5’-RDACGAGBACRTCTTCRGT-3’) – produces a 217 bp amplicon product. The reaction conditions in the first and the second round of the PCR consisted of the initial denaturation step at 94°C for 3 min followed by 35 cycles with denaturation at 94°C for 30 s, primer annealing at 50°C for 30 s and extension at 72°C for 1 min. The bar-coded PCR products from six *nod*A libraries (soil and nodule libraries from three plants species) were sequenced with a Roche 454 GS Junior (following the manufacturer’s protocols) generating an average of 3000-4000 reads per library. All obtained sequences were subjected to filtration by quality (quality score higher than 25), length (longer than 170 bp) and separating into libraries according to barcodes in QIIME. The sequencing data were deposited at the NCBI short read archive under the bioproject number PRJNA297503. We introduced the term “popset” to designate a set of *nod*A gene sequences from nodule or soil rhizobia population.

The multiple alignments of the remained sequences within each popset were performed with ClustalW as implemented in MEGA (Tamura et al. 2013). Sequences with frameshift errors were removed. The resultant multiple alignment for each popset did not contain gaps.

### *NFR5* amplification and sequencing

DNA from plant leaf pools was isolated by AxioPrep kit (Axigen) and was used as the template DNA for PCR amplifications. Approximately 0.9 kb DNA fragments encoding all three LysM domains of the plant receptor gene, *NFR5*, were amplified with the following pairs of primers: forward “NFR5-for4” (5’AAGTCTTGGTTGTTACTTGCC-3’) and reverse “NFR5-Grev3” (5’-CACCTGAAAGTAACTTATCYGCA-3’) for *Vicia sativa*; forward “NFR5-for4” and reverse “NFR5-Grev3” (5’-TGCAGTCTCAGCTAATGAAGTAC-3’) for *Lathyrus pratensis*; forward “NFR5-for4” and reverse “NFR5-Grev6” (5’-CATACATTGTTGGCTTGCTTAC-3’) for *Trifolium hybridum*. The standard PCR protocol was used: initial denaturation at 95°C for 3 min, 30 cycles with denaturation at 94°C for 30 s, primer annealing at 48°C for 30 s, extension at 72°C for 1 min and final extension for 4 min. PCR fragments were extracted from agarose gel ((Onishchuk et al. 2015)) and cloned into the plasmid pTZ57R/T (Thermo Scientific, Lithuania). For each plant species 100 randomly selected cloned fragments of *NFR5* genes were sequenced by Sanger method in an automated ABI 3500xL sequencer (Applied Biosystems) using standard M13 (−20) and (−26) primers. Sequences were deposited in the GenBank database under the PopSet accession number 1041522217. The multiple alignment of 100 sequences was performed with ClustalW as implemented in MEGA6(Tamura et al. 2013).

### Gene trees

The total *nod*A gene sequences from nodule and soil popsets aligned with ClustalW were clustered into operational taxonomic units (OTUs) at 95% nucleotide sequence identity threshold using the UCLUST algorithm implemented in QIIME 1.9. Neighbor-Joining (NJ) dendrogram based on the numbers of differences between representatives of each OTU was constructed in MEGA6 and rooted using the outgroup *nod*A gene sequence of *Sinorhizobium meliloti* (GenBank ID AZNW01000092.1).

### Diversity analysis

We quantified the diversity of haplotypes among rhizobia popsets using three indices: bias-corrected Chao1 index of the haplotype richness (Chao 1984), Simpson 1-D index of haplotype evenness (Simpson 1949) and Shannon H entropy index (Shannon 1948). Nucleotide diversity π was calculated as the mean difference between two randomly picked sequences in the popset.

In order to provide statistical support for differences between nodule and soil popsets, distributions of test statistics were obtained by independently subsampling with replacement of 2500 sequences from each popset in 2000 trials and bootstrapping of nucleotide positions 1000 times. All of the distributions passed the normality test (Kolmogorov-Smirnov test, p-value > 0.01). We tested the following null hypothesis for each diversity index: the diversity index value for a nodule popset was lower or equal than the diversity index value for the corresponding nodule popset. Welch’s t-test was used to evaluate the null hypothesis at 0.01 significance level.

To compare the levels of nucleotide diversity π between plant and nodule rhizobia popsets, we additionally constructed the distributions of π statistics for each plant population subsampling with replacement of 70 sequences in 2000 trials from each plant *NFR5* popset which initially contained 100 sequences. After that, we randomly formed pairs of π diversity values from the obtained distributions for the plants popsets and the respective nodule popsets. The total set of 6000 pairs (2000 per plant sp.) was taken as a paired sample data. The relationship between host plants (*NFR5* gene) and nodule rhizobial (*nod*A gene) population diversity was assessed as the value of the Spearman’s rank correlation coefficient for the paired sample data. The values higher than 0.8 were taken to imply monotonic relationship in π between plant and nodule rhizobial posets.

All calculations were implemented in MATLAB. The link to the GitHub repository containing MATLAB scripts for the diversity analysis is provided in the Supplementary file.

### Statistical methods for detecting selection

The dN/dS ratio is a widely used measure to quantify the selection pressure acting on a set of homologous protein-coding gene regions, where dN and dS are two measures of divergence between species, with dN corresponding to the number of nonsynonymous substitutions per non-synonymous site and dS to the number of synonymous substitutions per synonymous site. pN and pS statistics are analogous to dN and dS statistics but are used for levels of polymorphism within a population rather than divergence (Kryazhimskiy and Plotkin 2008).

Before calculating the pN/pS ratio, we analysed the independency of the pN and pS statistics. The analysis of linkage between nucleotide positions within the sequenced region of the *nod*A gene was performed within each joint popset (nodule + soil) as follows. First, for each pair of polymorphic nucleotide positions, we applied the chi-square test (with pooling) and extracted pairs of positions that were significantly non-independent (FDR p-value < 0.001). Second, for each position in the obtained pairs, we determined whether it was mostly a synonymous or non-synonymous polymorphic site (See Supplementary text). Third, we counted the number of sites that formed pairs with both synonymous and non-synonymous polymorphic sites and the total number of sites in linked pairs. We considered pN and pS to be independent of each other if the former number was twice as low as the latter number. Under the non-independence of pN and pS, the pS statistic does not reflect neutral mutations and the pN/pS ratio became inconsistent. Thus, we analysed pN and pS separately, mostly focusing on pN as natural selection acts on nonsynonymous changes. Comparing the nodule and soil popsets, we assumed that the increase of pN in one of them indicates the potential presence of stronger positive selection in it or the presence of stronger negative selection in the other popset. We hypothesized that the pN value in a soil rhizobial popset is higher of equal than the pN value in the respective nodule popset. We tested this hypothesis for each legume species separately performing Welch’s t-test considering 0.01 level of significance as described above.

Tajima’s D statistics represents the difference between the observed and the theoretically expected nucleotide diversity (Tajima 1989). If the mutations are neutral and the population adheres to the Wright-Fisher assumptions (Hartl and Clark 2007), Tajima’s D equals zero. Values significantly higher than zero indicate the deficit of rare alleles (e.g. due to a recent decrease in population size or balancing selection), and values significantly lower than zero indicate the excess of rare alleles (e.g. due to a recent population size expansion or purifying selection). In order to identify the selection type underlying the transformation of a soil popset to the respective nodule popset, we compared the values of Tajima’s D between the popsets. We considered the hypothesis that the Tajima’s D in the soil popset is higher or equal to that in nodule popset. We tested this hypothesis using Welch’s t-test at 0.01 level of significance.

The calculations were performed using MATLAB PGEToolbox (Population Genetics Evolution Toolbox). The link to the GitHub repository containing MATLAB scripts for the analysis of selection is provided in the Supplementary file.

### Topological organization of diversity in plant and rhizobia popsets

Let two populations of different sizes are represented by a set of aligned sequences. Let *p* be a population index, *p* ∈ {1,2}. At the first step, the method identifies the unique haplotypes and their frequencies in each population, denoted as 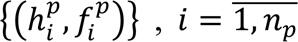, where *n*_*p*_ is the number of unique haplotypes in the population *p*. Let 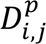 be the symmetric distance matrix between each pair of haplotypes in a population 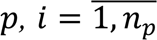. At the second step, the hierarchical agglomerative clustering method merges haplotypes into clusters until the number of clusters equals to predefined number *m*. Let fix a population *p* and omit this index. The clustering algorithms starts with placing each haplotype to its own cluster which is described by two parameters: frequency *f*_*i*_ and mean difference σ_*i*_ (initialized to zero). Then, the following procedure is repeated until *m* clusters is achieved. Two clusters, *i*′-th and *j*′-th with the smallest pairwise distance 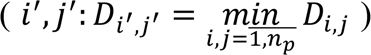 are merged into one new cluster. A distance from this cluster to a *k*-th cluster is calculated as follows: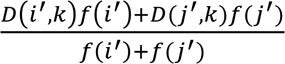 The frequency of the new cluster is *f* (*i)* + *f*(*j*) and the mean difference of the new cluster is *D*_*i*′_,_*j*′_.

The described hierarchical clustering is applied to each populations and yields the *m* clusters of haplotypes with the reduced distance matrix between them 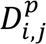, the frequencies 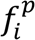 and the mean differences within clusters 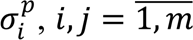 In order to normalize 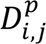 and 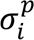 value between two populations we divided these values by the median across 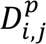. If a cluster contains only one haplotype and its 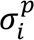 equals to zero, we set 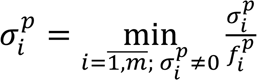.

At the third step, the set of clusters for each population was translated into *K*-dimensional Euclidian space by Metric multidimensional scaling (Metric MDS) that transforms the distance matrix 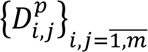 into a set of coordinates 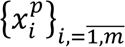. Then, the Gaussian mixture model (GMM) is introduced for the population *p* as follows:

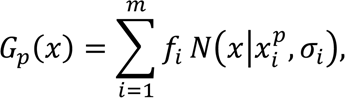

In the current project we took *K* = 3 as it was enough for low values of the stress function in MDS procedure.

The adjustment of two GMMs is carried out by Procrustes superimposition: the minimization of the dissimilarity between mixtures (Δ*G*) via translating, rotating and mirror reflection. The lower Δ*G* value is, the more similar two GMMs are and, consequently, more similar topologies of two population structures are.

For each plant species we performed two comparisons: “plant population vs. nodule popset” and “plant population vs. soil popset”. We joint rhizobial nodule and soil populations before clustering and MDS, and separated them before the Procrustes analysis. This manipulation ensured that the same haplotypes in both comparisons were taken into account in the same way. We tested the null hypothesis that the Δ*G* value in the first comparison is higher or equal to the Δ*G* value in the second comparison. In other words, the similarity between nodule rhizobia and plant popset topologies is not higher than the similarity between soil rhizobia and plant popset topologies. For each of the two comparisons, we obtained the set of Δ*G* values bootstrapping of sequences in rhizobial popsets. To test the hypothesis, we compared two obtained sets of Δ*G* values by one-sided Mann-Whitney U test with 0.01 level of significance.

For visual comparison of plant and rhizobial populations we constructed tanglegrams based on adjusted GMMs after Procrustes superimposition. A tanglegram is a diagram with a pair of two binary trees with matching leaves connected by edges. To construct it we built two NJ gene trees for *m* plant *NFR5* clusters and *m* rhizobium *nod*A clusters based on the between-cluster distance matrices and plotted two trees face to face. A pair of leaves from two trees was connected by an edge if a 3D point corresponded to one leave was located within the five closest points to a point of another leave and vice versa.

The link to the GitHub repository containing MATLAB scripts for topological beta-diversity analysis is provided in the Supplementary file.

## Results

### *nod*A gene: OTUs and cluster analysis

*Nod*A gene libraries (for root nodules and for soil samples) were sequenced by NGS technology. Sequencing and filtration of *nod*A gene libraries produced a total of 22463 sequences, for an average of 3750 sequences per popset. Clustering by 95% sequence identity produced 15 OTUs, which frequencies differed between the popsets. The NJ tree constructed for OTU representatives showed that the rhizobium population consisted of two clusters (orange and blue in Figure 2). Almost all sequences from the first cluster (9 OTUs) belonged to Vicia and Lathyrus popsets, while sequences from the second cluster (6 OTUs) were mostly detected in Trifolium popsets. The converse was also true: most of the Vicia-Lathyrus popset belonged to the first cluster (90% average), while most of the Trifolium popset belonged to the second cluster. This distribution of OTUs was in agreement with the “cross-inoculation groups” concept, which refers to the fact that separate groups of legume species can be successfully inoculated by only the specific groups of rhizobia. We attributed the first cluster to *R.leguminosarum* bv. *viciae* and the second to *R.leguminosarum* bv. *trifolii.* For further analysis, we kept in Vicia-Lathyrus popsets only sequences from the first cluster, and in Trifolium popsets, only sequences from the second cluster.

**Figure 2.**
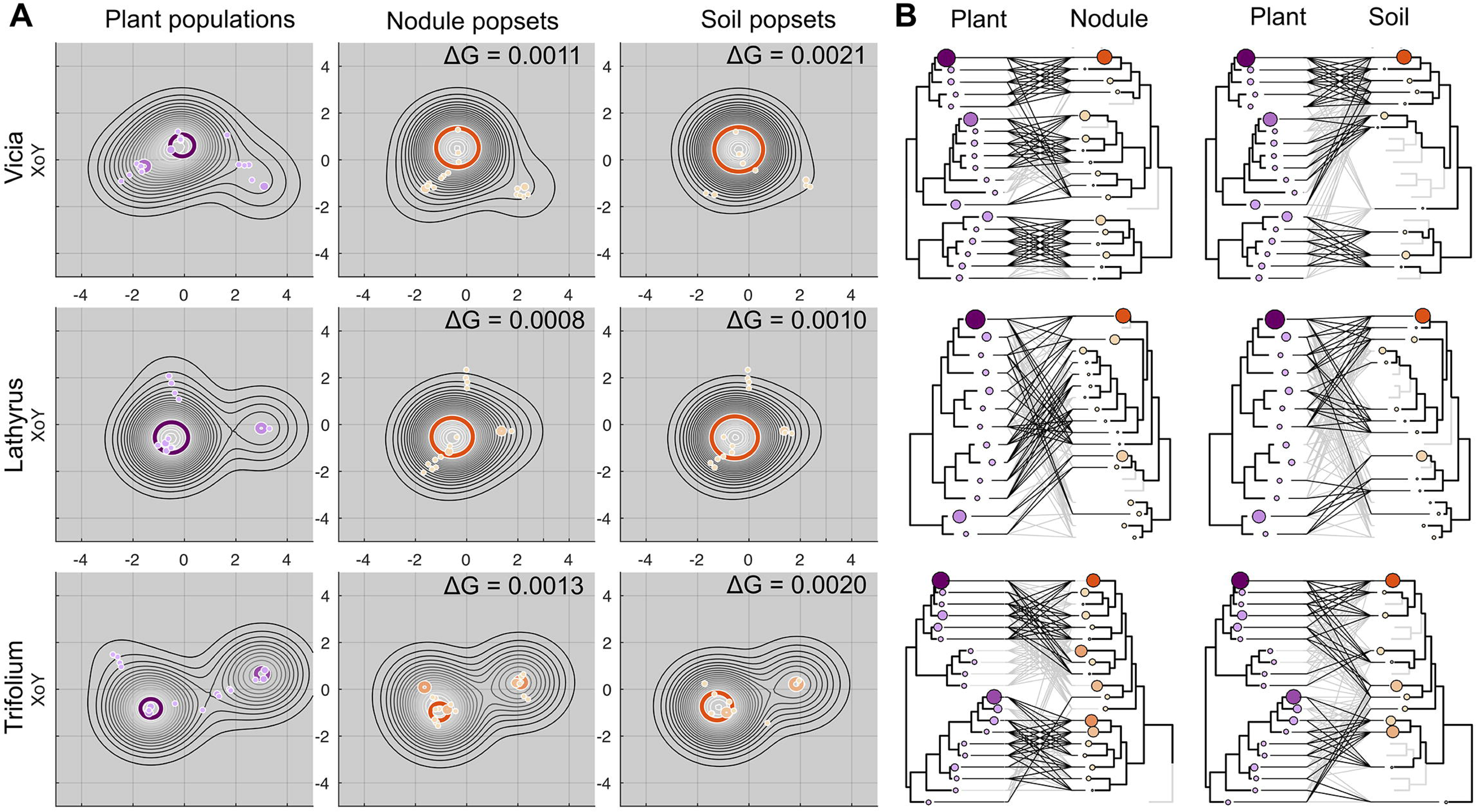
Clustering of *nod*A gene sequences after OTU-picking analysis. Columns correspond to six popsets (see text). Values in cells represent the numbers of OTU sequences in a popset, widths of rectangles reflect to the log of these values. The NJ tree of OTU-representatives forms two clades corresponding to *Rhizobium leguminosarum* biovars from different cross-inoculation groups: bv. *viciae* and bv. *trifolii*.

### Differences between soil and nodule populations of rhizobia

For each of the six *nod*A gene popsets, we examined the following measures of population diversity: number of haplotypes (Supplementary Figure S1), Chao1 index of richness, Simpson 1-D index of evenness, Shannon H index of entropy, and nucleotide diversity π. All diversity indexes are significantly higher in nodule popsets compared to soil popsets (Figure 3.A-D; Welch’s t-test, p<0.01).

### Analysis of selection

Analysis of population structures revealed the significant linkage between synonymous and non-synonymous polymorphic sites, suggesting that the pN and pS statistics were not independent (see Supplementary Text). Further comparison of pN/pS values between nodule and soil popsets did not establish the significant difference (Supplementary Fig. S7). Despite the inconsistency of pN/pS in our case, we observed that the values of both statistics, pN and pS, were significantly increased (Welch’s t-test, p-values < 0.01) in the nodule popsets, indicating that both non-synonymous and synonymous diversity was elevated there (Table 1). The values of Tajima’s D were significantly lower than 0 in all of the rhizobial *nod*A popsets, indicating the presence of negative selection (Table 1). However, within each plant, they were significantly higher in nodule popsets than in the soil popsets (p-values < 0.01), suggestive of relaxation of negative selection or admixture of balancing selection in the former. Trifolium popsets displayed the highest values of this statistic, consistently with the strongest admixture of balancing selection, while the Lathyrus popsets, the lowest.

**Table 1.** Values of pN, pS, pN/pS and Tajima’s D statistics for the popsets of different origin. The difference in pN/pS values between nodule and soil popsets for each plant was not significant. The significance (p-value < 0.01) of the difference in values between nodule and soil population for each plant are marked with “**”.

| Popset |  | pN | pS | pN/pS | Tajima's D |
| --- | --- | --- | --- | --- | --- |
| Vicia | nodule | 0.0170 | 0.0228 | 0.7450 | -2.2644 |
|  | soil | 0.0075 ** | 0.0099 | 0.7600 | -2.5219 ** |
| Lathyrus | nodule | 0.0065 | 0.0086 | 0.7570 | -2.5609 |
|  | soil | 0.0051 ** | 0.0072 | 0.7186 | -2.5977 * |
| Trifolium | nodule | 0.0245 | 0.0335 | 0.7301 | -2.0294 |
|  | soil | 0.0167 ** | 0.0232 | 0.7184 | -2.2496 ** |

### Relationship between the bacteria and host plant diversities

We calculated the diversity levels π in each plant population and found that the ranking of the three species was the same as the ranking based on the π nucleotide diversity values in corresponding nodule popsets. By bootstrapping the plant and rhizobial nodule popsets we estimated the Spearman correlation between these popsets’s nucleotide diversities. The 0.89 value, that was higher than the predefined threshold indicated that the monotonic relationship between diversities in popsets of plant *NFR5* gene sequences and in bacterial *nod*A nodule popsets is statistically significant (Figure 4).

### Concordance of gene trees

A visual comparison of topologies of plant *NFR5* trees with those of corresponding rhizobial *nod*A trees for nodule and soil popsets revealed that the topology of clades in the plant trees was more similar to the topology of clades in the nodule trees than in the soil trees (Supplementary Figure S3). Based on this observation, we proposed that the gene tree of the nodule rhizobia is more similar to that of the host plant than the gene tree of the soil rhizobia.

To formally test this, we developed a method for comparing structures (topologies) between two popsets. We tested the null hypothesis that the topological similarity between a nodule rhizobia popset and a plant popset is not higher than the topological similarity between the soil rhizobia popset and the plant popset. This hypothesis was rejected at the 0.01 level of significance. We constructed tanglegrams that also illustrated a higher similarity of the topology of the nodule rhizobial *nod*A gene trees (Figure 5, left tanglegram) than the soil rhizobial *nod*A gene trees (Figure 5, right tanglegram) to that of plant NFR5 gene tree.

## Discussion

Symbiotic interactions represent a special case of ecological interactions when one of the partners provides an “environment” for another. In “On the Origin of Species” (Darwin 1872), Charles Darwin proposed that “the life of each species depends in a more important manner on the presence of other already defined organic forms, than on climate”. This is particularly true for organisms in deeply integrated symbiotic systems.

Here, we traced the coordination in levels of population diversities between partners within the essential components of the rhizobium-legume signalling system: plant symbiotic receptor gene *NFR5* and *Rhizobium* symbiotic gene *nod*A involved in the synthesis of signalling molecules Nod factors, ensuring the first stage of partner recognition (Oldroyd 2013). The matching was detected in three phenomena. The first is the increase of population diversity levels in nodule rhizobial population (mutualist phase) in a comparison with soil rhizobial popsets (free-living phase). The second is the monotonic relationship between the host plant diversity and the nodule rhizobial diversity. The third is the similarity in topological diversity between host plant population and nodule rhizobial population. All of these observations illustrated the aspect of symbiotic interactions, which we refer to as the Evolutionary Moulding effect. Under this effect the soil rhizobial population *nod*A gene pool “is moulded into rigid matrix” of host-plant population *NFR5* gene pool forming nodule pool of *nod*A and ensuring (1) the sufficient increase of the diversity level in nodule pool to meet the host plant needs (2) the same trend in the matching diversity levels between rhizobial and host-plant across legume species (3) the “topological” matching of *NFR5* with *nod*A gene trees between host-plant popset with nodule rhizobial popset, but not with soil rhizobial popset.

The increased diversity (evenness, richness and nucleotide polymorphism) in nodule popsets in comparison with respective soil popsets seem unexpected while only 90 nodules formed each nodule popset. Indeed, as the formation of each nodule popset involves a bottleneck of only ∼ 90 nodules the nodule population can be expected to have a reduced population size and therefore, reduced diversity. However, if the nodule population was formed non randomly, this increase is not surprising. There are several possible explanations to the observed trends: (i) selection imposed by plants on soil populations which leads to increased diversity in nodule populations; (ii) increased mutation rate in bacterial populations under stress conditions inside root nodules (Krasinikov and Melkumova 1963; Roumiantseva et al. 2004); (iii) negative selection in the soil which reduces diversity there. The last explanation is unlikely, as the selection pressure in the soil hardly affects the symbiotic genes. The second explanation remained only hypothetical as it was not well described and had no much attention in recent studies (Krasinikov and Melkumova 1963; Roumiantseva et al. 2004). By exclusion, the first explanation is the most likely. The analysis of the selection imposed by plants revealed significantly increased nonsynonymous diversity (pN) and Tajima’s D values in the nodule popsets. This may be indicative of weaker negative selection in a nodule popset in a comparison with the respective soil popset but is also consistent with a contribution of balancing selection or presence of stronger population structure in the former. Earlier it was shown that in symbiotic systems, besides the above-mentioned types of selection, negative frequency-dependent selection in favour of rare genotypes during the competition of rhizospheric bacteria for root nodulation(Amarger and Lobreau 1982) may also play an important role (Andronov et al. 2015; Provorov and Vorobyov 2000; Provorov and Vorobyov 2006).

The pronounced influence of plant on the formation of nodule population was demonstrated in the monotonic correspondence between diversity in plant popsets and diversity in respective nodule rhizobial popsets (Spearman correlation = 0.89). Another convincing evidence of the plant-imposed effect was obtained with the specially developed method to compute “topological beta-diversity” – difference in topological structures of two population sets (plant and rhizobia) of sequences. The similarity between *NFR5* and *nod*A gene trees of plant and nodule popsets was significantly higher than the same between plant and soil popsets. This result demonstrated the transformation of the initial soil *nod*A pool by the template of the host plant receptor pool. Our conclusion is in line with the numerous works studying the interplay between diversities of host plant and rhizobia (Paffetti et al. 1998; Andronov et al. 2003; Bena et al. 2005; Bailly et al. 2006; Depret and Laguerre 2008; Rangin et al. 2008; Barrett et al. 2016; Vuong, Thrall, and Barrett 2016; Österman et al. 2011), however, in most cases we cannot directly compare these studies due to the differences in the experimental design.

The observed similarity between nodule rhizobial *nod*A gene popsets and plant *NFR5* receptor gene popsets revealed the hierarchical organization of effective interaction: two symbionts should be genetically compatible at the single organism level and also at the population level. The process of forming this interaction could be explained metaphorically as the Evolutionary Moulding: shaping the population structure of one symbiont using the population structure of another symbiont as a “matrix”. The important point in this shaping is the difference between evolutionary rates in plants and bacteria. The bacteria have a significantly higher evolutionary rate than plants, therefore the diversity of *nod*A gene in bacterial populations, like the flexible genetic material in the Evolutionary Moulding, reflected the shape of more “rigid” diversity of *NFR5* receptor gene in plant populations. We hypothesise that under the Evolutionary Moulding effect two symbiotic populations tend to relax the incoordination of genetic diversities between two parts of the symbiont-host signalling system, that is mostly achieved by a faster evolving partner. In the current project the faster evolving partner is rhizobia, however, we assume that if plants hypothetically had evolved with higher rates than bacteria, we would have expected that the rhizobial component would have played a role of the matrix (or mould) in terms of the Evolutionary Moulding. It should be also highlighted, that under the Evolutionary Moulding, the transformation of the “more flexible” symbiont population does not necessarily lead to its increased population diversity. When a population of a “more rigid” symbiont is, for example, homogeneous (conservative in polymorphism), the population diversity of the flexible symbiont could decrease.

According to the Evolutionary Moulding effect, the relationship in population diversity between rhizobia and host-plant may be observed not only within the pair of *nod*A-*NFR5* genes (which are related through the Nod factor) but also within any pair of interplaying genes from plant and bacterial sides. We propose that genome-wide searching of “matching” genes under the Evolutionary Moulding can be an extension to the traditional methods of functional analysis of genes.

At present, we can only hypothesize the molecular mechanism of the Evolutionary Moulding in rhizobium-legume symbiosis. We suppose that polymorphism in symbiotic genes of a rhizobial population is probably associated with the structural diversity of Nod factors produced by this population, in particular with variations in the unsaturated fatty acid tail. Nowadays, an impressive progress is achieved in the resolution and accuracy of the methods to analyse the Nod factor structure (Poinsot et al. 2016) and its docking to the receptor proteins (Malkov et al. 2016). We believe that such approaches will facilitate the analysis of Nod factor structural variation produced by rhizobial populations and, as a result, the molecular mechanisms in the Evolutionary Moulding.

**Figure 3.** (**A-D)**: Diversity measures. Letters “V”, “L”, “T” denote Vicia, Lathyrus, Trifolium popsets respectively. Letters “Soil” and “Node” denote soil and nodule popsets. Precise formulas to compute diversity measures are shown in Supplementary TableS1. Mean values of diversity measures are presented in the Supplementary TableS2.

**Figure 4.** The monotonic relationship between the π diversity levels in nodule popsets and plant popsets. Dots represent the distribution of π obtained by bootstrapping.

**Figure 5.** Comparison of plant popsets with the bacteria popsets from nodules and soil. **(A)**: Projections of Gaussian mixture models for three plant *NFR5* popsets and six rhizobial *nod*A popsets after the Procrustes analysis on the XoY plane (see Supplementary Figures S4-S6 for other projections). The values of Δ*G* correspond to the difference between the GMMs for plant and rhizobial (nodule or soil) popsets; differences between Δ*G* values in each row are significant (p<0.05). A visual comparison of projection confirms this trend. For example, in the “Vicia” row the rhizobium nodule popset has two peaks that remind two peaks in the plant popset and more distinct than in the rhizobium soil popset **(B)**: Tanglegrams for each plant species: between *NFR5* population and rhizobial *nod*A populations from nodule (left tanglegrams) and soil (right tanglegrams).

## Acknowledgements

This study is supported by RSF (Russian Science Foundation) grant 14-26-00094(P).

