## Supplementary Materials for "The Evolutionary Moulding in plant-microbial symbiosis: matching population diversity of rhizobial *nod*A and legume *NFR5* genes"

The link to the GitHub repository containing MATLAB scripts is https://github.com/iganna/popselection.git

### Supplementary tables

**Table S1.** Measures of diversity

| | |
|---|---|
| Bias-corrected Chao1 index (richness): | $$S_{Chao1} = n + \frac{n_1(n_1 - 1)}{2(n_2 + 1)},$$ where $n$ is the total number of observed sequences, $n_1$ is the number of singletons and $n_2$ is the number of doubletons. |
| Simpson $1 - D$ index (evenness): | $$S_{Simpson} = 1 - D,$$ where D is calculated as follows: $$D = \frac{\sum_{i=1}^{n_h} n_i(n-1)}{n(n-1)}$$ $n$ is the total number of observed sequences, $n_h$ is the number of unique haplotypes, $n_i$ is the number of sequences corresponding to the $i$-th haplotype. |
| Shannon H index (entropy): | $$H_{Shannon} = -\sum_{i=1}^{n_h} p_i \ln(p_i),$$ where $n_h$ is the number of unique haplotypes, $p_i$ is the proportion of sequences corresponding to the $i$-th haplotype. |
| $\pi$ diversity: | $$\pi = 2\sum_{i=1}^{n_h}\sum_{j=1}^{i-1} p_i p_j \pi_{i,j},$$ where $\pi_{i,j}$ is the number of differences per site between $i$-th haplotype and $j$-th haplotypes. |

**Table S2.** Mean values of four diversity measures in rhizobial popsets. The significance (p-value < 0.01) of the difference in values between nodule and soil population for each plant are marked with “**”.

| Popset | | Chao1 (x1e3) | Simpson | Shannon | Nucleotide diversity |
|---|---|---|---|---|---|
| Vicia | nodule | 1.0016 | 0.6246 | 2.6204 | 0.0198 |
| | soil | 0.6398** | 0.5420** | 2.1015** | 0.0087** |
| Lathyrus | nodule | 0.7198 | 0.5958 | 2.2994 | 0.0076 |
| | soil | 0.7093 | 0.5024** | 2.0344** | 0.0061** |
| Trifolium | nodule | 1.6829 | 0.9000 | 3.7762 | 0.0283 |
| | soil | 0.9526** | 0.8042** | 3.0526** | 0.0197** |

### Supplementary figures

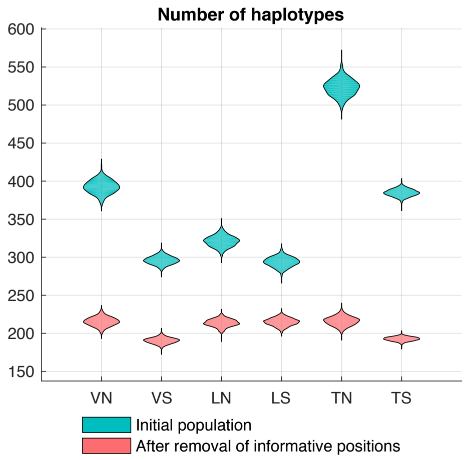


**Supplementary Figure S1.** Number of haplotypes within rhizobial popsets. Letters “V”, “L”, “T” denote Vicia, Lathyrus, Trifolium popsets respectively. Letters “N” and “S” denote nodule and soil popsets.

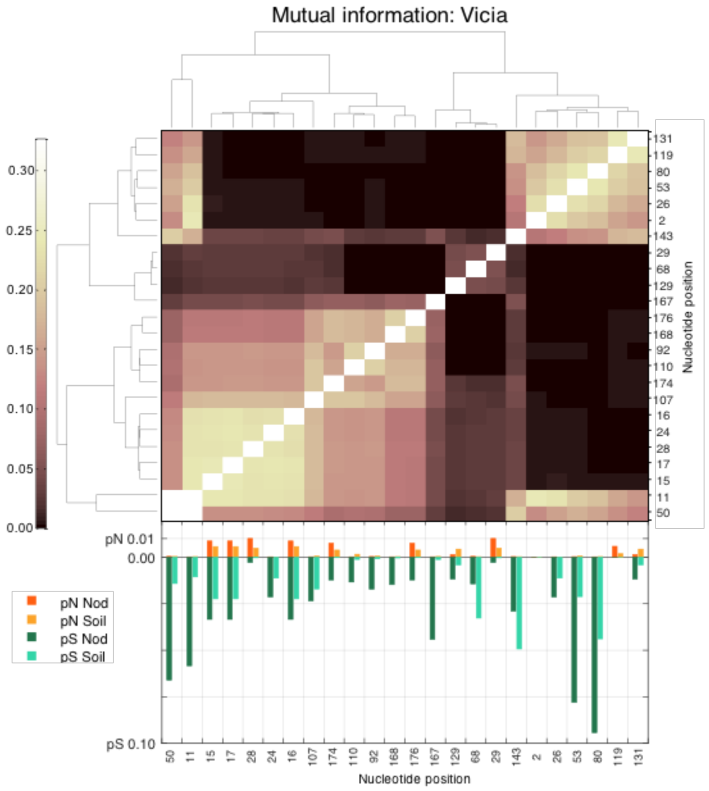


**Supplementary Figure S2.** Mutual information between linked nucleotide positions within *nod*A region in Vicia joint (nodule+soil) popset. Positions form two clusters according to the UPGMA algorithm, cityblock distance. **Bottom figure**: The pN, pS values in codons containing linked positions was calculated separately for nodule (darker colour) and soil (brighter colour) popsets.

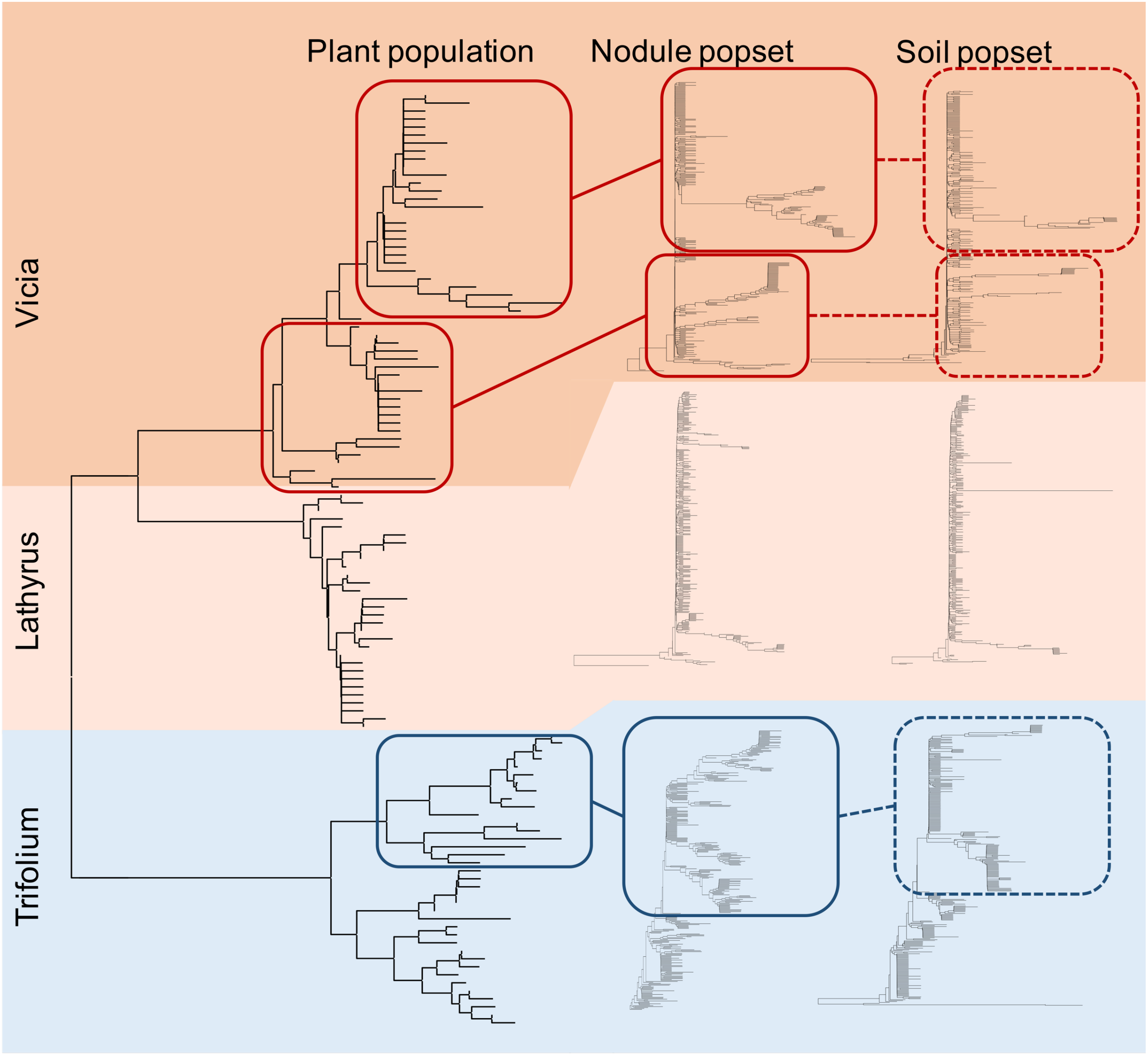


**Supplementary Figure S3.** Neighbour-joining (NJ) tree built for unique three plant haplotypes simultaneously contains three the major clades and is presented on the left. NJ for nodule and soil popsets was extracted from the NJ trees for joint (nodule+soil) popsets with preservation tree topology, therefore, they are visually comparable. All the trees were rooted using outgroup sequences that were removed before visualisation.

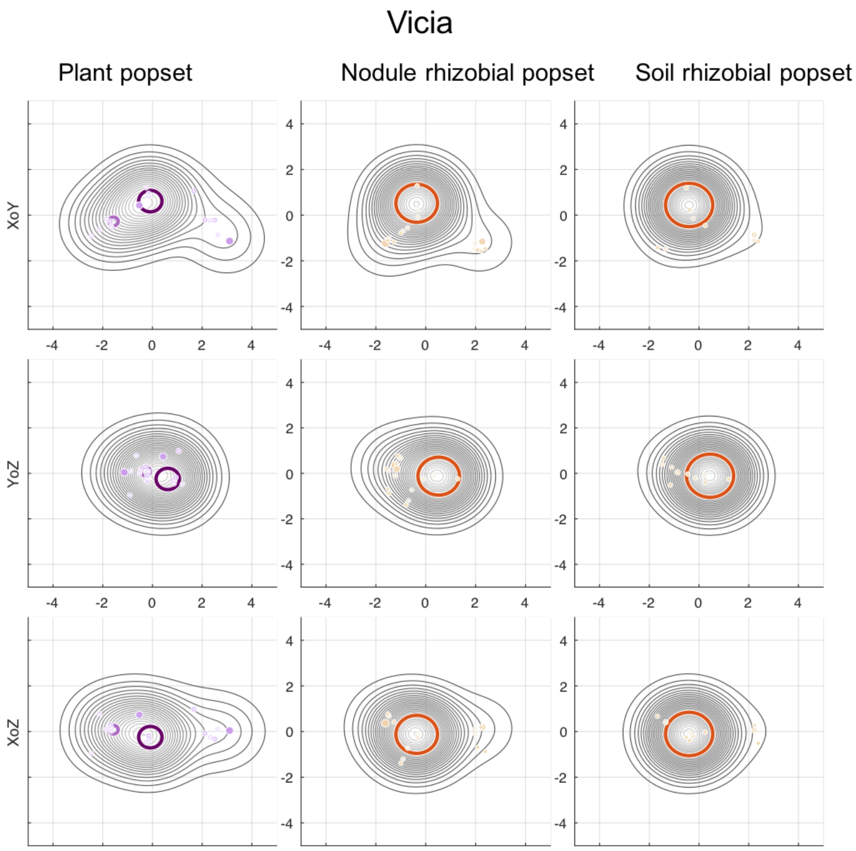


**Supplementary Figure S4.** Three projections of the Gaussian mixture models for Vicia host-plant popset and Vicia rhizobial popsets (nodule and soil)

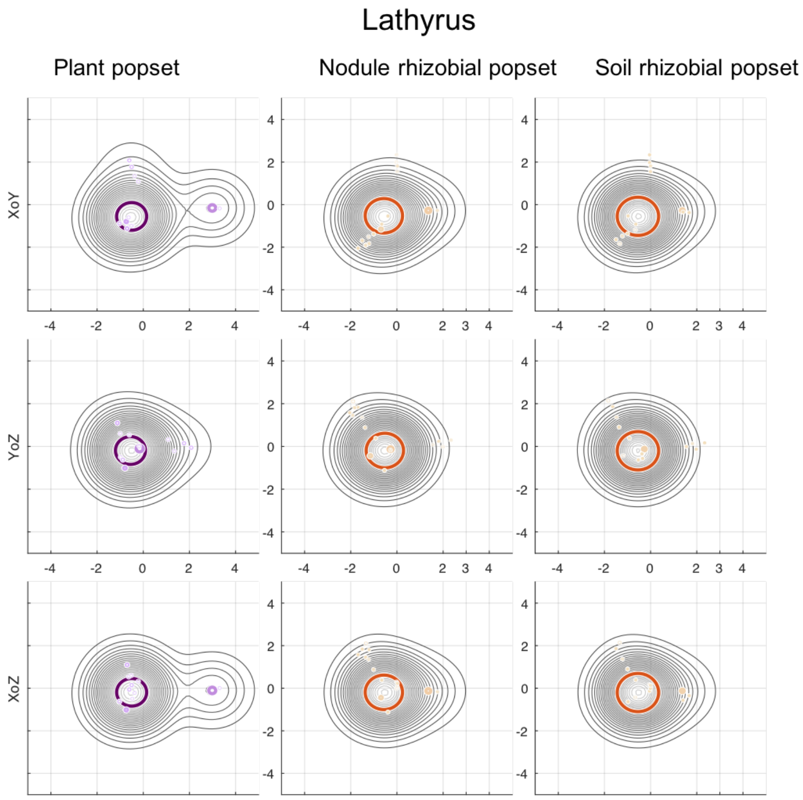


**Supplementary Figure S5.** Three projections of the Gaussian mixture models for Lathyrus host-plant popset and Lathyrus rhizobial popsets (nodule and soil).

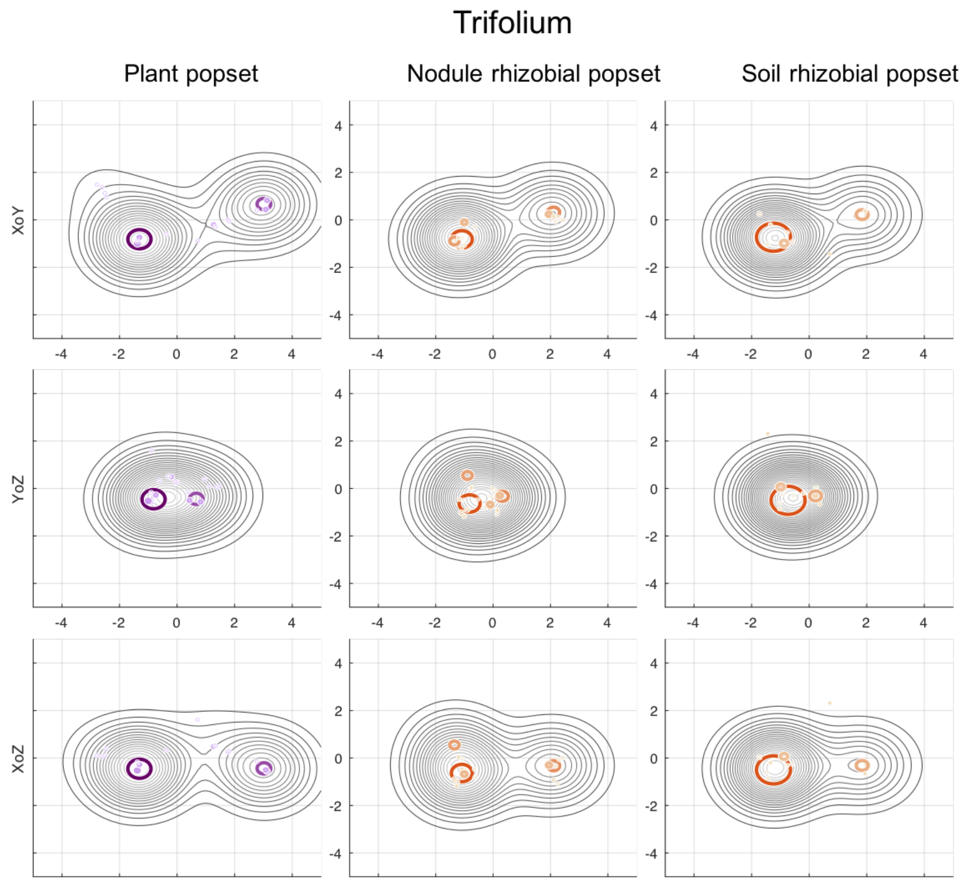


**Supplementary Figure S6.** Three projections of the Gaussian mixture models for Trifolium host-plant popset and Trifolium rhizobial popsets (nodule and soil).

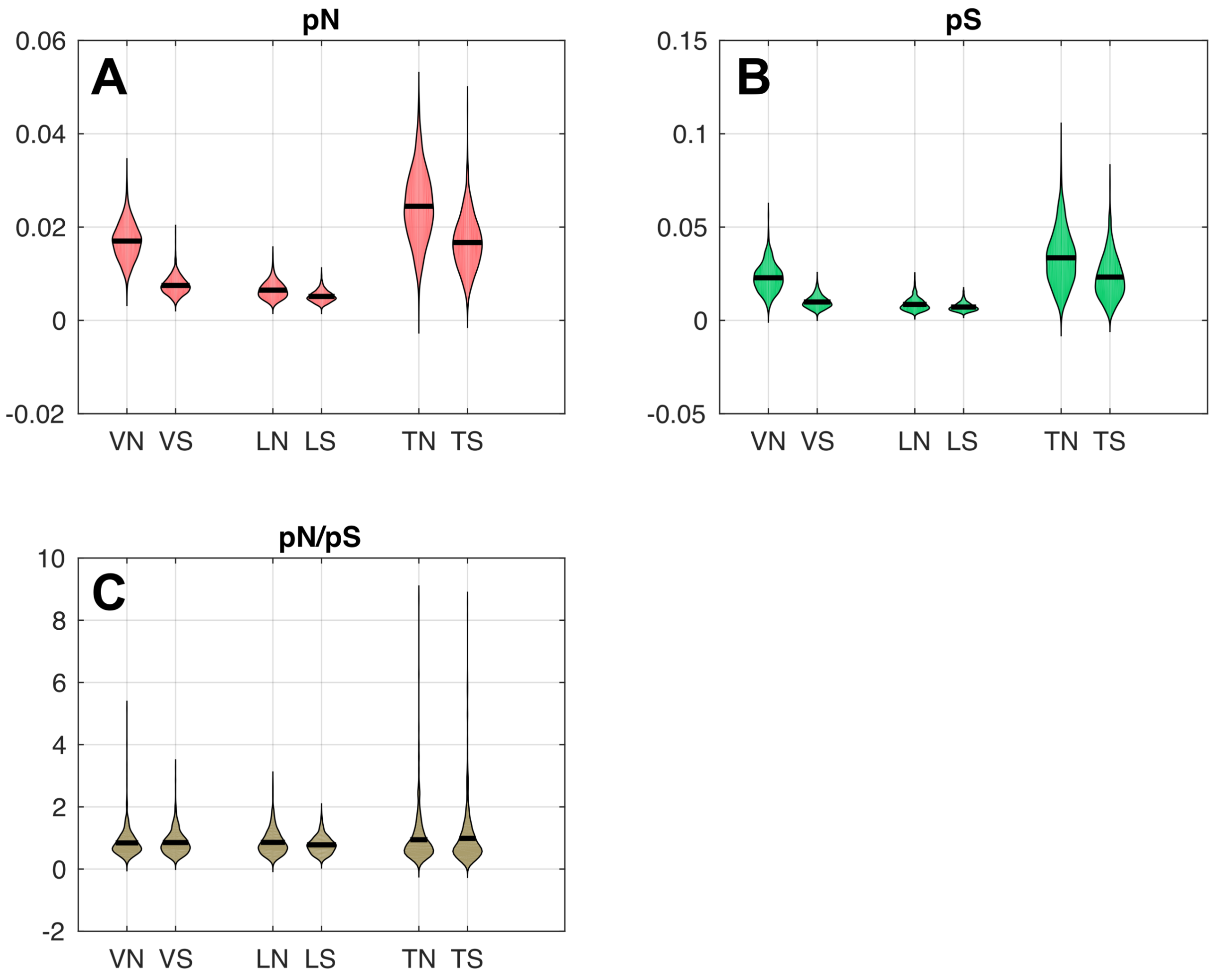


**Supplementary Figure S7.** Distributions of pN and pS statistics values. **(A,B)** Differences between in pN and pS values between nodule(N) and soil(S) popsets were significant (p-value < 0.01). **(C)** The difference in pN/pS values was not detected (p-value > 0.01). Letters “V”, “L”, “T” denote Vicia, Lathyrus, Trifolium popsets respectively. Letters “N” and “S” denote nodule and soil popsets.

### Supplementary Text

#### Construction of gene trees for host-plant and rhizobial popsets

Detailed *nod*A gene trees for popsets were obtained in three steps. For each plant species, we first combined nodule and soil popsets and identified unique haplotypes in the joint set of sequences. Next, we constructed a NJ dendrogram based on p-distances between all unique haplotypes and rooted it as above. Finally, we extracted two subtrees from the dendrogram according to the sequences from nodule and soil popsets. This algorithm yields structures of nodule and soil popsets that could be easily compared. Outgroups were removed from each tree before visualisation. The phylogenetic tree for *NFR5* gene sequences from all three plant species was also constructed using NJ algorithm based on p-distances.

#### Linked positions in *nod*A regions

Within *nod*A region joint popsets (nodule+soil) for each legume species we tested all pairs of nucleotide positions to be significantly linked. We detected 24, 22 and 19 positions forming significantly linked pairs (chi-square test with pooling, BH adjusted p-value = 0.001) within Vicia, Lathyrus and Trifolium joint popsets, respectively. Overlapping between some linked pairs evinced the groups of linked positions that were confirmed after biclustering the symmetric matrix of mutual information between the positions (Fig. 4).

We calculated pN and pS statistics at each codon containing detected positions and analysed the composition of identified groups. All groups of associated substitutions contained both non-synonymous and synonymous substitutions, to be more specific, each position with high pS in the corresponding codon placed in one group with at least one position with high pN in the corresponding codon. Moreover, there are several positions with high pN belonged to one group. The groups of linked positions did not place on separate regions of *nod*A fragment, conversely, the nucleotide areas of the groups overlapped. When we compared sets of linked positions with the corresponding sets of informative positions we found them 98% overlapped.

**Analysis of possible PCR-related effects**

To test whether the observed difference could be a result of underestimation of diversity in the soil due to difference in the starting quantity of DNA templates between nodule and soil DNA pools, we analysed the number of nucleotide sited influencing to the increase of diversity in nodule popset comparing to the soil popset. At the 0.01 level of significance we reject the influence of experimental conditions responsible for this increase.

To detect whether the difference in the starting quantity of DNA templates between nodule and soil DNA pools was responsible for the low diversity in soil popset, we analysed the number of the nucleotide positions that contributed the most to diversity level. For this purpose, we ranked all of the nucleotide positions in the region of *nod*A gene according to the Shannon's entropy calculated in each nucleotide position within a joint popset (nodule + soil corresponding to one legume species). The higher rank corresponded to the higher value of Shannon's entropy. We successively removed (without return) one nucleotide positions with the highest rank, recalculated distributions of diversity indexes (Chao1, Simpson $1 - D$, Shannon H and $\pi$ diversity) for nodule and soil popset and applied Welch's t-test to test the null – diversity within soil popset is higher or equal than within nodule popset - as described above. While the hull hypothesis was rejected, we continued the process. We counted the number of positions that were removed before the null was not rejected and denoted them as influencing positions. Assuming the presence of experimental artefacts leading to underestimation of soil diversity, influencing positions involved the most of the polymorphic positions, therefore the probability that a polymorphic position was an influencing one, should be higher than 0.5. We set the one-sided hypothesis that one category (influencing positions) is more frequently occur than other category (non-influencing positions). The respective one-sided Binomial test (with the probability parameter $p = 0.5$ and the number of traits equals to the number of polymorphic sites) was performed to test the null hypothesis at 0.01 level of significance.

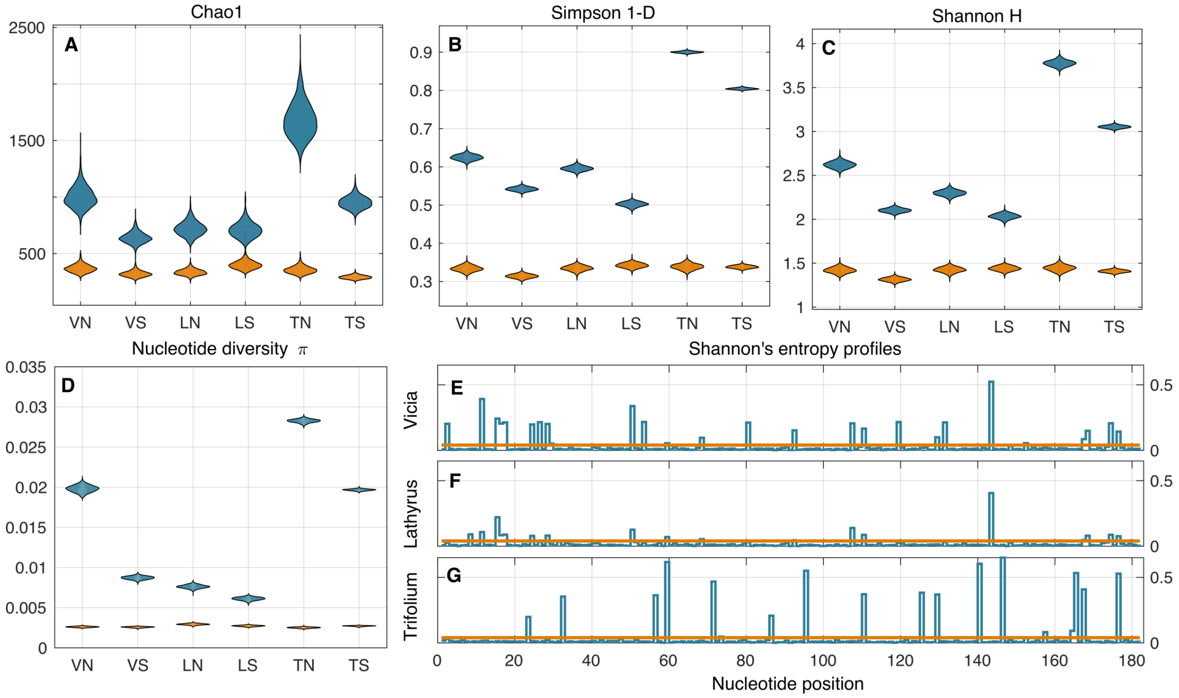


**Figure** (**A-D)**: Diversity measures. Letters "V", "L", "T" denote Vicia, Lathyrus, Trifolium popsets respectively. Letters "N" and "S" denote nodule and soil popsets. For example, "VN" means the Vicia nodule popset. Blue violins represent diversity levels in initial popsets; orange violins represent diversity levels in popsets after the removal of informative positions. (**E-G)**: Shannon's entropy profiles for *nod*A gene region for three legume species. The orange threshold (0.4) separates informative positions (peaks) and non-informative ones.
